# Histone H1x in mouse ventral hippocampus correlates with, but does not cause behavioral adaptations to stress

**DOI:** 10.1101/2023.11.06.565881

**Authors:** R. Kijoon Kim, Natalie L. Truby, Gabriella M. Silva, Joseph A. Picone, Cary S. Miller, Rachael L. Neve, Xiaohong Cui, Peter J. Hamilton

## Abstract

Prior research has identified differential protein expression levels of linker histone H1x within the ventral hippocampus (vHipp) of stress-susceptible versus stress-resilient mice. These mice are behaviorally classified based on their divergent responses to chronic social stress. Here, we sought to determine whether elevated vHipp H1x protein levels directly contribute to these diverging behavioral adaptations to stress. First, we demonstrate that stress-susceptible mice uniquely express elevated vHipp H1x protein levels following chronic stress. Given that linker histones coordinate heterochromatin compaction, we hypothesize that elevated levels of H1x in the vHipp may impede pro-resilience transcriptional adaptations and prevent development of the resilient phenotype following social stress. To test this, 8–10-week-old male C57BL/6J mice were randomly assigned to stressed and unstressed groups undergoing 10 days of chronic social defeat stress (CSDS) or single housing respectively. Following CSDS, mice were classified as susceptible versus resilient based on their social interaction behaviors. We synthesized a viral overexpression (OE) vector for H1x and transduced experimental mice with either H1x or control GFP within vHipp. Following viral delivery, we conducted social, anxiety-like, and memory-reliant behavior tests on distinct cohorts of mice. We found no behavioral adaptations following H1x OE compared to GFP controls in susceptible, resilient, or unstressed mice. In sum, although we confirm vHipp protein levels of H1x correlate with susceptibility to social stress, we observe no significant behavioral consequence of H1x OE. Thus, we conclude elevated levels of H1x are correlated with, but are not singularly sufficient to drive development of behavioral adaptations to stress.

## Introduction

Chronic stress is a risk factor for developing neuropsychiatric syndromes including post-traumatic stress disorder and major depressive disorder^1^. Yet, not all individuals that experience chronic stress develop these syndromes. The neurobiological correlates that govern vulnerability to developing behavioral maladaptations to chronic stress remain incompletely elucidated, and a more complete understanding may inform the development of novel therapeutics^2,3^. Animal models of stress experience offer a useful research method to interrogate the brain molecular and epigenetic mechanisms that drive the diversity of stress-induced behavioral phenotypes^4^.

The Chronic Social Defeat Stress (CSDS) paradigm is a widely utilized method to subject laboratory rodents to social stress^5^. Studies applying CSDS in mice have identified and interrogated many differences in brain epigenetic, transcription, and protein-level responses associated with individual variability in stress^6–8^. Conventionally, stress-induced behavioral phenotypes are stratified into stress ‘susceptible’ versus ‘resilient’, where resilience is defined as the capacity to endure chronic social stress without developing detectable deficits in social interaction (SI) and susceptibility is defined as the development of reduced SI behaviors following CSDS. The application of transcriptomics and metabolomics has demonstrated that the resilient phenotype is a biologically active state involving brain neuroadaptations characterized by a transcriptional and metabolic state distinct from unstressed or susceptible mice, as well as identified a wide range of potential target genes that specifically orchestrate an individual animal’s resilience to stress^7–10^.

Many recent studies have demonstrated roles of core histone protein variants and post-translational modifications (PTMs) in coordinating stress response, memory, and fear conditioning. H1 variants and PTMs are known to respond to external stimuli and regulate chromatin compaction^11–13^, but the role of neuronal H1 variants in regulating behavioral adaptations to stress is not yet fully understood. Previous multiomic analysis of male mice post-CSDS found histone variant H1x to be differentially regulated within the ventral hippocampus (vHipp) between susceptible, resilient, and unstressed mice^8^, a brain region known to encode stress and anxiety-related learned behaviors^14–16^. Further, it is known that H1x induces the strongest chromatin compaction of all H1 variants^12^, a chromatin conformation associated with transcriptional silencing. Thus, we hypothesize that heightened levels of H1x within the mouse vHipp, either by natural or viral overexpression (OE) mechanisms, would impede the expression of resilience-specific molecular adaptations and result in the emergence of susceptible-like behaviors following chronic stress experience.

Here, we present a comprehensive behavioral assessment of viral OE of H1x within vHipp. We developed a novel H1x OE vector and validated intra-vHipp delivery and efficacy using western blot and immunohistochemistry methods. Further, we confirm intra-vHipp OE of H1x does not induce behavioral changes in stress-naïve mice, leading us to hypothesize H1x requires an interaction with stress experience in order to control the emergence of the susceptible phenotype. However, upon post-CSDS behavioral testing, we instead observed H1x OE did not alter social or anxiety-like behaviors in any condition. Collectively, this work demonstrates H1x, a protein that was strongly correlated with susceptibility to social stress, does not play a causative role in determining an animal’s susceptibility versus resilience to social stress. These data provide important additional context for the biological mechanisms that do, and do not, drive development of stress-induced behavioral adaptations and point to a more nuanced relationship between neuronal gene expression and organismal behaviors.

## Methods

### Subjects

Male C57BL/6J mice (8 to 10 weeks old) from Jackson Laboratories were used. Mice were group housed (5 mice/cage) on a 12-hour light/dark cycle (lights on at 6am/off at 6pm) with ad libitum access to food and water. All mice were used in accordance with protocols approved by the Institutional Care and Use Committees at Virginia Commonwealth University School of Medicine.

### Viral packaging

We sub-cloned the *H1fx* gene into HSV expression plasmids via ThermoFisher Scientific gateway LR Clonase II cloning reaction and Gateway LR Clonase II Enzyme mix kit (catalog number 11791-020 and 11971-100). Colonies were Maxi-prepped (Qiagen Cat # 12163) and shipped to the Gene Delivery Technology Core at Massachusetts General Hospital for HSV packaging. Once packaged, aliquots were made and stored in -80°C to be used in viral gene transfer through stereotaxic surgery.

### Viral gene transfer

Stereotaxic surgeries targeting the vHipp were performed as previously described^17,18^. Mice were anesthetized with I.P. injection of ketamine (100 mg/kg) and xylazine (10 mg/kg) dissolved in sterile saline solution. Mice were then placed in a small-animal stereotaxic device (Kopf Instruments) and the skull surface was exposed. 33-gauge needles (Hamilton) were utilized to infuse 1.0 μL of virus at a rate of 0.2 μL/minute followed by a 5 minute rest period to prevent backflow. The following coordinates were used to target the vHipp: Bregma: anterior-posterior: - 3.4 mm, medial-lateral +3.2 mm, dorsal-ventral -4.3 mm, 3° angle.

### Histology

3 days following viral gene transfer, when GFP expression remained readily detectable, mice were transcardially perfused with 0.1 M sodium phosphate buffer, followed by 4% paraformaldehyde (PFA) in 0.1 M phosphate buffer. Brains were removed and postfixed in 4% PFA overnight at 4°C (Brains: n = 5 for HSV-GFP, n = 7 for HSV-H1x). Following postfix, whole brains were cryoprotected via submersion in 15% sucrose in PBS with 0.05% sodium azide at 4°C for 24 hours followed by 30% sucrose in PBS with 0.05% sodium azide at 4°C for at least 24 hours and until sectioning. Coronal sections at 20 μm thickness containing the vHipp were sliced on a Leica CM1860 cryostat and mounted on Superfrost™ white slides.

### Immunohistochemistry

Following cryosectioning, tissue slices underwent antigen retrieval in boiling Tris/EDTA buffer at pH 8.0 for 20 minutes. Sections were then permeabilized with 0.1% Triton X-100 in TBS for 10 minutes and blocked with 10% goat serum and 1% BSA in TBS for 2 hours at room temperature. Slides were incubated at 4°C overnight with rabbit anti-H1x primary antibody (Abcam ab31972) diluted 1:800 in 1% BSA in TBS, then with goat anti-rabbit Alexa Fluor 594 (mCherry) secondary antibody (Abcam ab150080) diluted 1:1000 in 1% BSA in TBS at room temperature. Additionally, a GFP booster stain (Chromotek Alexa Fluor 488) at 1:500 and DAPI stain (Thermofisher D1306) at 1:1000 dilution was included with the secondary antibody. Sections were mounted using ProLong Gold Antifade and coverslipped. Slides were kept at 4°C in a light-blocking slide box until imaging. Z-stack images of each section were taken at 20x on a Keyence BZ-X810 microscope (n = 5-8 images per brain).

### Image Analysis

Image analysis was completed using ImageJ. First, maximum intensity projections of each Z-stack were calculated, and the image was split between GFP, mCherry, and DAPI channels. Cell nuclei were identified using trainable Weka segmentation^19^, and the resulting image was converted to a region of interest mask. This mask of cell nuclei was applied to the mCherry channel image. Integrated Density within the nuclei regions was calculated and compared between viral treatment groups using GraphPad Prism 10.

### Chronic Social Defeat Stress

CD1 retired breeder mice were screened for aggressive behavior as previously described^5^. A C57BL/6J mouse is subjected to 10 consecutive days of 10-minute social defeats. In each defeat session, a C57BL/6J mouse is placed in the home cage of a single-housed CD1 mouse, where the CD1 mouse attacks the intruding C57BL/6J mouse. Following defeat, the mice are separated by a clear plastic perforated divider overnight. C57BL/6J mice are antagonized by a novel CD1 aggressor for each day of defeat. For accelerated social defeats (Supp. Fig. 3), these defeat sessions were completed twice per day for 5 days separated by 6 hours.

### Behavioral Testing

Behavioral analyses were performed automatically by video tracking software (Ethovision Noldus)^20^. All behavioral tests were performed in a specified behavioral suite under red light.

### Social Interaction

Behavioral testing was performed 24 hours after the final social defeat. Stressed mice are single housed for the duration of behavior testing. SI was performed as previously described^5^. C57BL/6J mice were placed into the open arena (43×43×43cm) with an empty wire cage (10×5×30cm) at one side (interaction zone). Experimental C57BL/6J mice (45 mice per virus treatment across two cohorts) were given a 2.5 minute habituation session to explore the arena and then removed from the open arena. A novel CD1 aggressor was then placed within the wire cage (interaction zone), and the experimental mouse was placed back into the open arena for another 2.5 minute session. The arena was thoroughly cleaned with 70% EtOH before a new C57BL/6J mouse was placed in the arena and before the first trial, but not between sessions of an individual mouse’s test. Data was analyzed as time spent in the interaction zone without the aggressor compared to time spent within the interaction zone with the aggressor present. The data was then calculated into SI ratios by dividing the time spent in the interaction zone with the target mouse present by the time spent in the interaction zone (IZ) with the target mouse absent (SI ratio = time in IZ target present / time in IZ target absent.) Total distance moved (locomotion) and time spent exploring the center of the open field (thigmotaxis) was recorded and evaluated when the CD1 target was absent.

### Elevated Plus Maze

The EPM apparatus is constructed of black Plexiglas and consists of two open arms (33×6cm) and two closed arms (33×9.5× 20cm) facing connected by a central platform (5×7cm)^21^. The maze was kept elevated 63cm above the floor. C57BL/6J mice (45 mice across two cohorts) was placed individually in the right-side closed arm facing the center of the plus-maze. Placement of all four paws into an arm was registered as an entry in the respective arm. The time spent in each arm was recorded during the 5 min EPM test. The platform of the maze was cleaned with 70% EtOH following each trial and before the first trial. The percentage of time spent in the open arms was calculated (time spent in open arms/300 s * 100 = % time spent in open arms).

### Novelty Suppressed Feeding

Mice (cohort of 30) were food restricted for 24 hrs prior to NSF. The floor of an open field arena (43×43×43cm) was covered in bedding, and a single piece of highly palatable food was placed in the center of the open field. Experimental mice were then placed into the corner of the arena, and the time to approach and eat the food was recorded. Once the mouse approached and began to feed, or after 10 minutes if the mouse did not begin to feed, it was removed and returned to its home cage, where the latency to feed was again recorded. A latency of 10 minutes (600s) was recorded for mice that did not approach or eat the food. The bedding was changed between trials for different experimental mice.

### Fear Potentiated Startle (FPS)

Following established protocols^22^, mice (cohort of 30) were placed in sound insulated chambers, and baseline acoustic startle responses were recorded in millivolts by an accelerometer in the floor of the startle chamber (Day 1). Following 10 pairings of cue light to a foot shock (Day 2), their acoustic startle response with cue light on was compared to baseline startle (Day 3). Proportional startle responses were calculated as previously described^23^. To determine whether mice acquired a conditioned startle response, a one-sample t-test comparing proportional startle response to a theoretical mean of 0 was used. This value represents identical startle response with and without the cue light, which would correlate with a mouse that did not acquire fear conditioning. The worst learner defined as the least positive proportional startle response after conditioning from each virus treatment group was removed from the analysis. Mice then underwent 60 extinction trials over two days to only the cue light (days 4-5), and their startle response with and without the cue light was recorded again (day 6). Startle chambers are cleaned thoroughly with 0.1% quaternary ammonium chloride disinfectant between mice.

### Statistical Analysis

All data was analyzed in GraphPad Prism 10. Results were expressed as mean ± standard error and were analyzed parametrically. Groups were analyzed with Student’s two-tailed t-tests (Fig. 2B, C, Supp. Fig. 3B, C), one-way analysis of variance (Fig. 1B, C, Supp. Fig. 4), two-way analysis of variance (Fig. 3B-E, Supp. Fig. 1, Supp. Fig. 2, Supp. Fig. 5), or three-way analysis of variance (Fig. 4B-F, Fig. 5B, C) with Bonferroni’s post-hoc test. *P* < 0.05 was considered statistically significant.

**Figure 1:**
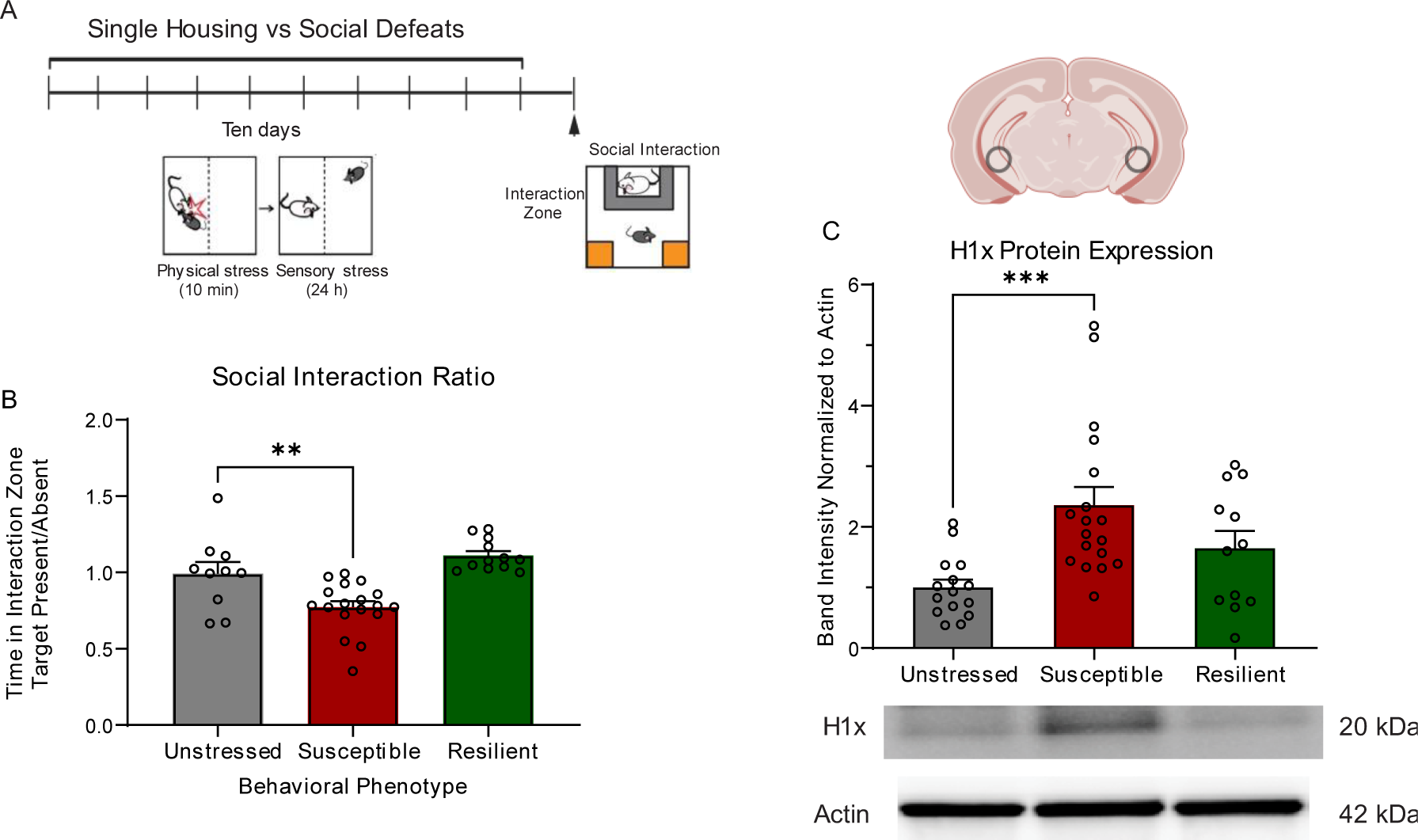
Stress-susceptible mice show increased hippocampal H1x protein expression following chronic social defeat. (A) Experimental timeline detailing 10-day chronic social defeat stress paradigm. (B) Stress-susceptible mice (n = 18) show significant decrease in social interaction ratio compared to unstressed mice (n = 10) and stress-resilient (n = 12) mice. (One-way ANOVA with Bonferroni’s correction, **: p = 0.0053) (C) Western blot analysis shows ventral hippocampus tissue from susceptible brains express significantly higher H1x protein levels compared to unstressed control brains, while resilient brains do not. (One-way ANOVA with Bonferroni’s correction, ***: p = 0.0010)

**Figure 2:**
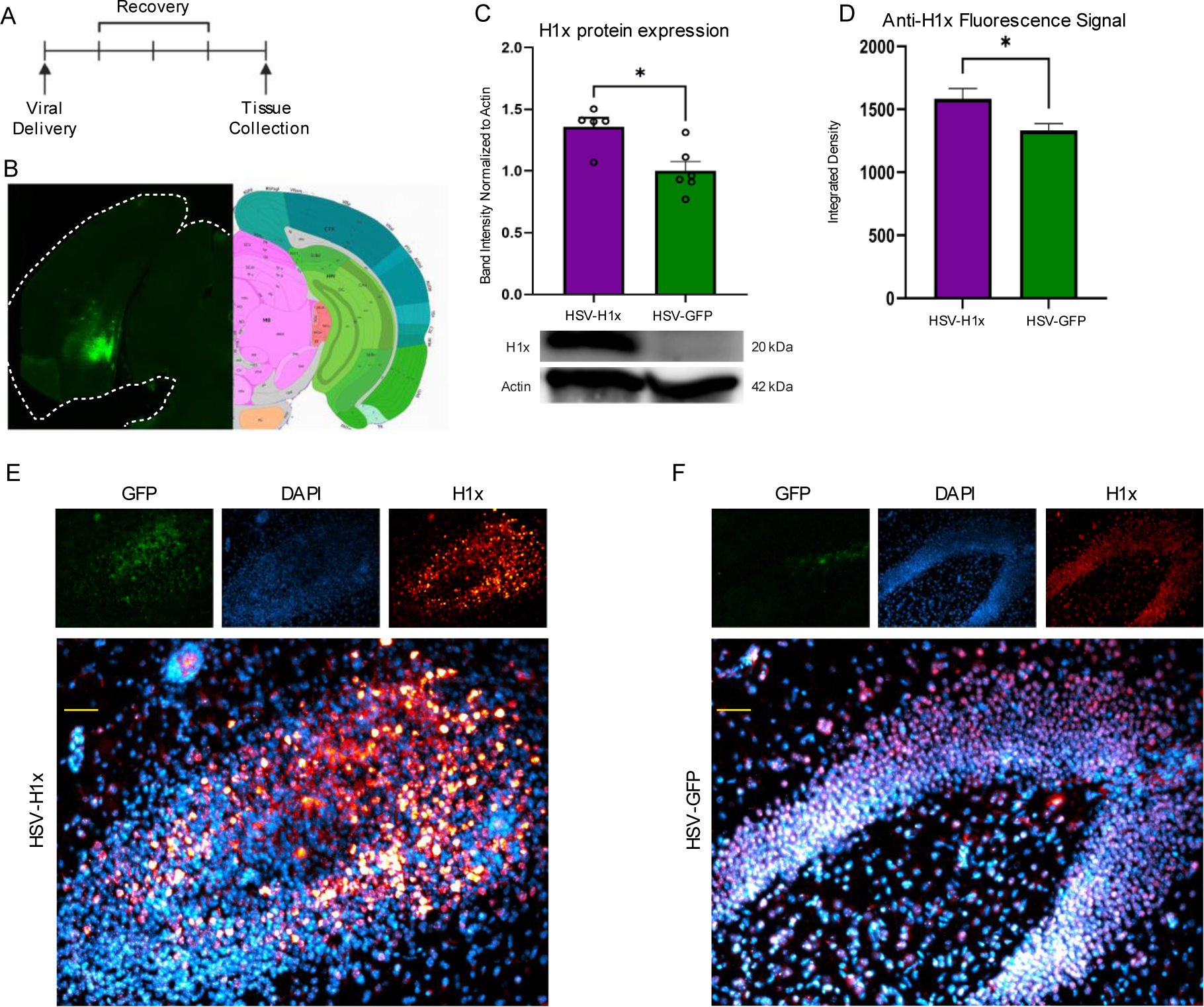
Intra-ventral hippocampus delivery of HSV-H1x induces overexpression of H1x specifically within cell nuclei. (A) Experimental timeline. Tissue for molecular assays was collected 3 days post-viral transduction. (B) Histologic analysis demonstrates appropriate delivery of virus to the ventral hippocampus, with the most intense GFP signal localized to the dentate gyrus. (C) HSV-H1x induces significant overexpression of H1x protein in ventral hippocampus compared to HSV-GFP as measured by western blot. (n = 5-6 per viral condition, Two-tailed t-test, *: p = 0.0193). (D) HSV-H1x induced significant increase in intra-nuclear H1x fluorescence signal compared to HSV-GFP (n = 5-7 brains per virus, 5-7 sections per brain, Two-tailed t-test, *: p = 0.0169). This spatial pattern of expression matches what would be required for H1x function under unmanipulated conditions. Fluorescence signal was quantified as integrated density within DAPI-positive nuclei identified by trainable Weka segmentation. (E-F) Representative images demonstrating increased fluorescence signal from H1x localized specifically within cell nuclei. GFP signal is not shown in the merged image for clearer visualization of H1x and DAPI colocalization, but was noted to be present in all images. (Scale bar = 50 µm).

**Figure 3:**
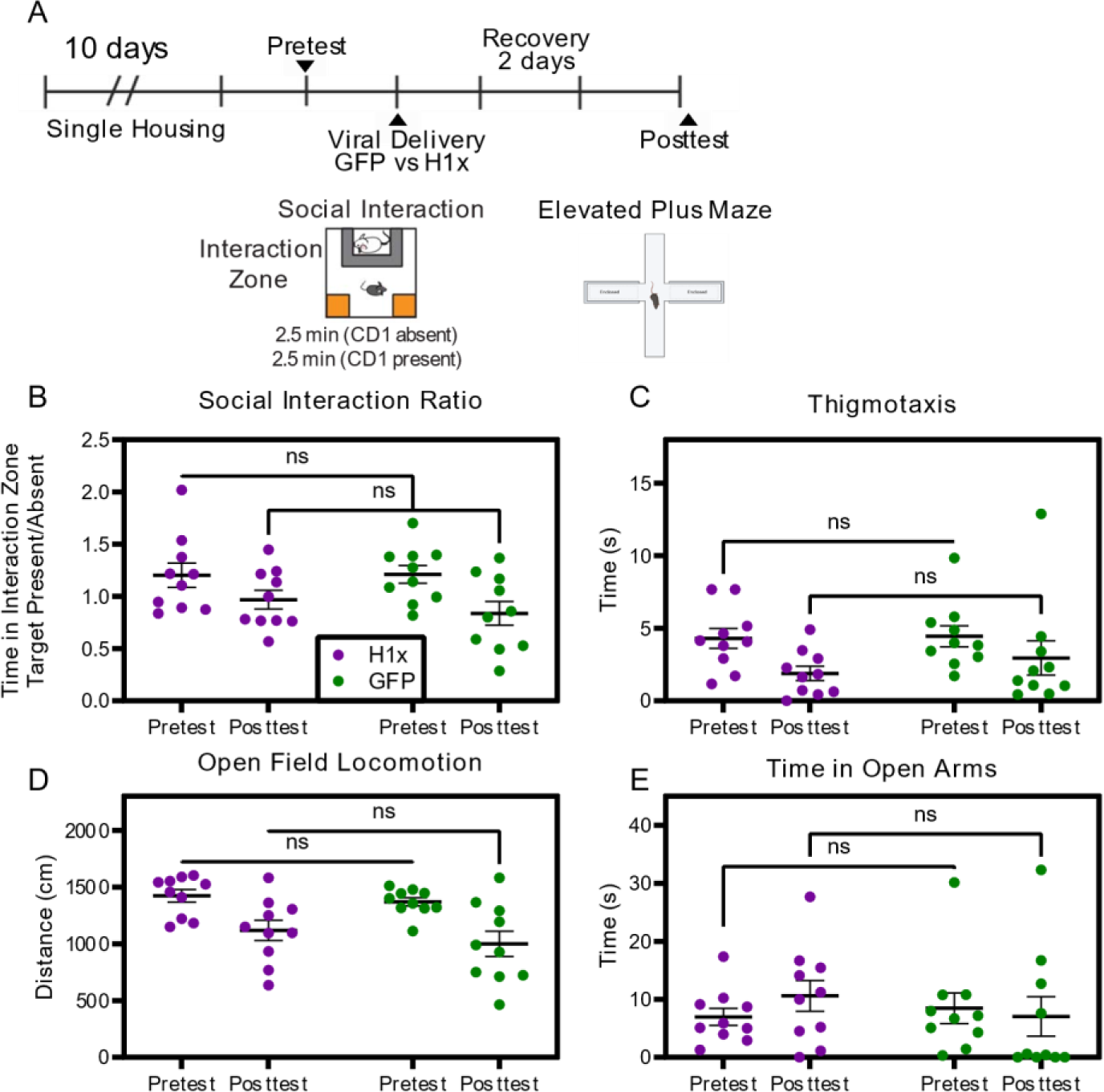
Intra-hippocampal overexpression of H1x in stress-naïve mice does not induce behavioral changes. (A) Experimental timeline including 10-day single housing experience. Pretest and posttest involved social interaction and elevated plus maze testing. Mouse populations receiving HSV-H1x versus HSV-GFP were balanced based on pretest behavior results. (B-E) No difference in behavior metrics in social interaction, open field, or elevated plus maze was observed between virus conditions (Two-way ANOVA with Bonferroni’s correction, p > 0.05).

**Figure 4:**
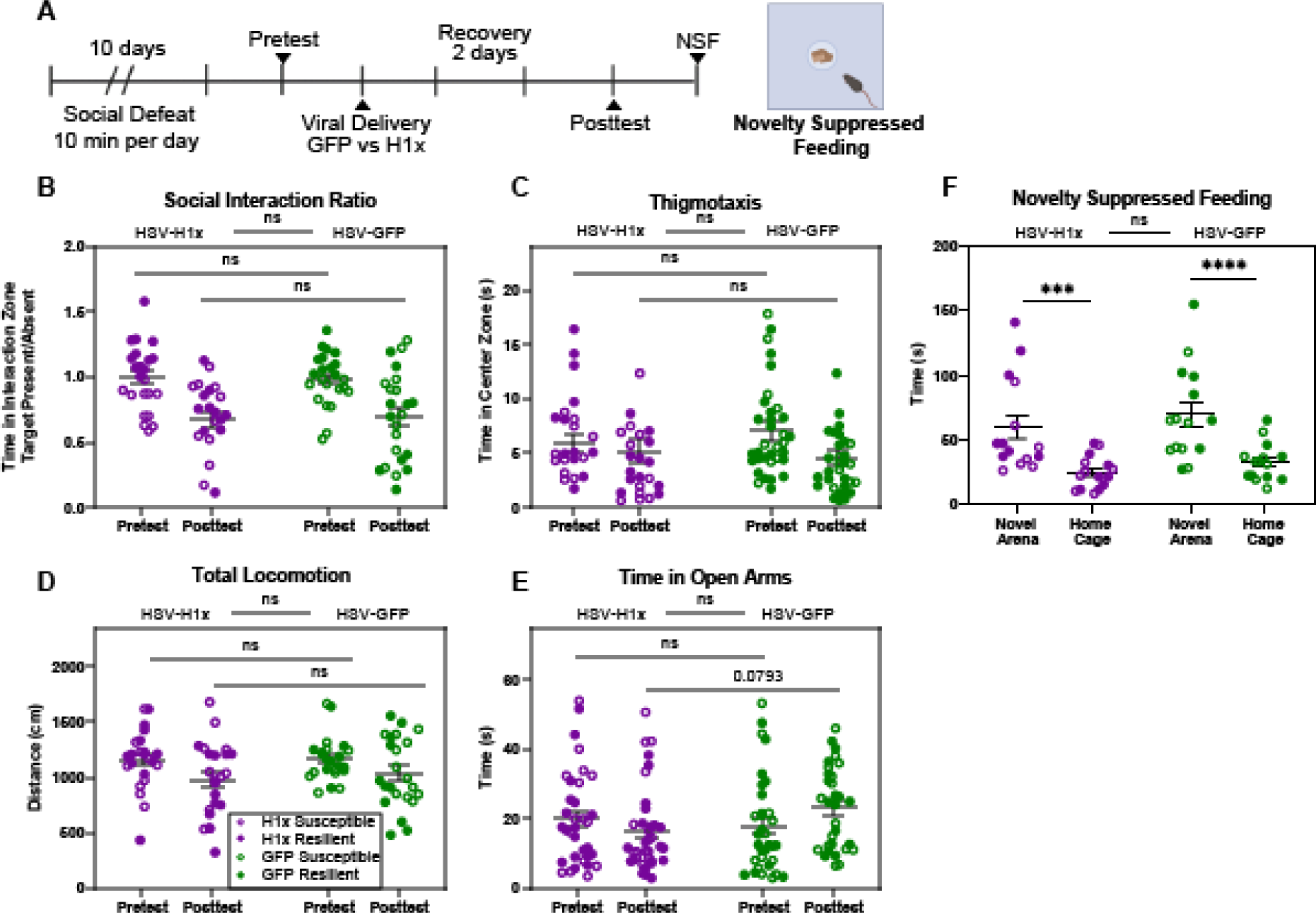
Intra-hippocampal overexpression of H1x does not cause development of behavioral deficits following social defeat stress. (A) Experimental timeline detailing 10-day chronic social defeat stress. Pretest and posttest involved social interaction and elevated plus maze testing. Mouse populations receiving HSV-H1x versus HSV-GFP were balanced based on pretest behavior results. (B-D) No behavior differences in social interaction or open field behaviors were found between HSV-H1x versus HSV-GFP treatment on the posttest (Three-way ANOVA with Bonferroni’s correction, p > 0.05). (E) Elevated plus maze data showed a significant test day by virus treatment interaction term (Three-way ANOVA with Bonferroni’s correction, p = 0.0491), however there was not a significant difference between groups on posttest (p > 0.05). When data was consolidated by viral treatment ignoring pretest social interaction phenotype, there was a trend (p = 0.0793) suggesting HSV-H1x mice explore the plus maze less on posttest than HSV-GFP mice. (F) Novelty suppressed feeding did not reveal differences between virus treatments. Mice treated with HSV-H1x showed similar latency to feed to HSV-GFP mice in both the novel arena and the home cage (p > 0.05). All mice showed a much shorter latency to feed in the home cage compared to the novel arena (HSV-H1x: p = 0.0003, HSV-GFP: p = 0.0001). Open circles denote susceptible mice, whereas filled circles denote resilient mice.

**Figure 5:**
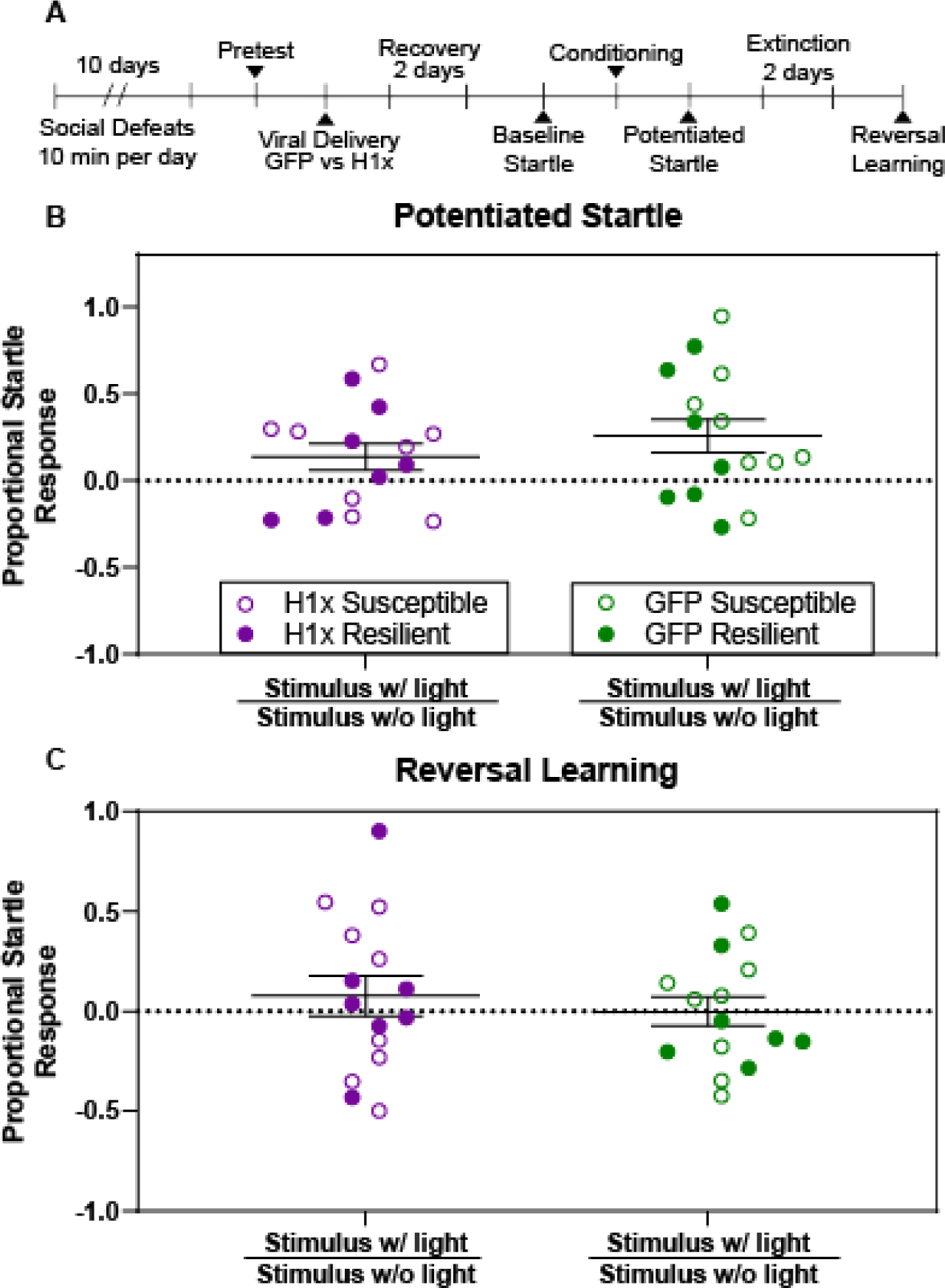
Intra-ventral hippocampus overexpression of H1x does not alter acquisition or reversal of fear potentiated startle. (A) Experimental timeline. Pretest involved social interaction testing. Mouse populations receiving HSV-H1x versus HSV-GFP were balanced based on pretest behavior results. (B-C) Both HSV-H1x and HSV-GFP treated mice acquire a potentiated startle response following 10 foot shock-cue light pairings (One-sample t-test, p = 0.0510 and 0.0076 respectively), and both groups successfully inhibit this startle response following 60 cue light only extinction trials (One-sample t-test, p = 0.3259 and 0.8164 respectively). Pretest social interaction phenotype was not found to significantly contribute to observed variance, so viral treatment groups were merged regardless of pretest phenotype (Three-way ANOVA, main effects p > 0.05). No significant difference was found between viral treatment groups in either potentiated startle or reversal learning sessions (Two-way ANOVA with Bonferroni’s correction, p = 0.6100 and 0.9611 respectively).

## Results

### Stress-susceptible mice express elevated levels of H1x protein within ventral hippocampus

We first sought to elucidate the vHipp protein expression patterns of H1x in the phenotypically distinct groups following chronic stress. To this end, we employed the CSDS model in male C57BL/6J mice and generated cohorts of susceptible, resilient, and unstressed control mice (Fig. 1A-B). In line with existing data^5^, we observed susceptible mice interact with CD1 target mice significantly less than unstressed or resilient mice (Fig. 1B). Following behavior testing, we performed a targeted western blot on microdissected vHipp to quantify H1x protein levels in this brain region (Fig. 1C). We discovered susceptible mice expressed significantly elevated protein levels of H1x compared to unstressed mice (p = 0.0007). Conversely, resilient brains expressed similar levels of H1x to unstressed controls (p > 0.05). This result demonstrates a correlation between elevated H1x expression and susceptibility to stress, in agreement with earlier proteomics data^8^. Yet, the question whether this difference in H1x expression is sufficient to drive manifestation of the susceptible behavioral phenotype remains unanswered. Thus, we aimed to probe the causative role of H1x in stress-induced behavioral adaptations by performing a comprehensive behavioral assessment following virally-mediated manipulation of H1x expression within the vHipp.

### Intra-ventral hippocampus viral delivery of H1x significantly increases H1x protein levels within the transduced nuclei

In order to assess the behavioral consequences of elevated H1x, we synthesized a bicistronic plasmid to express both H1x and a GFP reporter, and packaged the construct in herpes simplex virus (HSV-H1x). We also created a control virus containing only the GFP reporter for use as a non-functional control (HSV-GFP). Using stereotaxic surgery techniques, we demonstrated accurate and repeatable viral delivery to the vHipp (Fig. 2A-B). Next, we validated the expression efficacy of our viral vectors using a targeted western blot on microdissected vHipp tissue from transduced brains (Fig. 2C). As expected, vHipp tissue transduced with HSV-H1x expressed significantly higher levels of H1x protein compared to HSV-GFP control tissue (p = 0.0092). To determine whether increased H1x density is localized to the transduced neuron’s nucleus, as would be required for appropriate H1x molecular functionality, we performed immunohistochemical staining to confirm localization of virally overexpressed H1x within nuclei of transduced vHipp tissue. H1x signal colocalized with DAPI nuclear staining, without additional expression outside of nuclei (Fig. 2E-F). This provided further confirmation that our HSV-H1x virus overexpresses H1x protein without disrupting the intracellular distribution of H1x. We utilized trainable Weka segmentation to measure anti-H1x fluorescence signal within cell nuclei (Fig. 2D). In agreement with our western blot data, this approach found significantly increased anti-H1x signal within HSV-H1x brains compared to HSV-GFP brains (p = 0.0169). These viral validation assays give confidence in the efficacy of our HSV-H1x OE vector.

### Intra-vHipp overexpression of H1x in stress-naïve mice does not alter behavior

We next aimed to determine the behavioral consequences of intra-vHipp H1x OE. As H1x was determined to be uniquely correlated with stress susceptibility, we first posed the question whether H1x OE itself produces any behavioral changes in anxiety or social phenotypes. Here, we delivered HSV-H1x or HSV-GFP to the vHipp of stress-naïve mice and subjected them to SI and elevated plus maze (EPM) tests (Fig. 3A). We elected to utilize a pretest/posttest paradigm to ensure each cohort of mice receiving HSV-H1x and HSV-GFP respectively were behaviorally matched pre-surgery, allowing us to determine whether transduction with each virus altered a mouse’s behavior from its baseline.

We hypothesized that H1x would counteract stress-induced vHipp molecular adaptations and therefore exert no effect on the behaviors of stress-naïve mice. In line with these expectations, we did not observe any significant differences between mice treated intra-vHipp with HSV-H1x versus HSV-GFP on either SI or EPM posttest (Fig. 3B-E). Both HSV-H1x- and HSV-GFP-treated mice exhibited equivalent social interaction ratio, thigmotaxis, total locomotion, and time spent in open arms (p > 0.05). This indicates that H1x does not impact any of our tested behaviors in the absence of stress experience.

### Overexpression of H1x following social defeat stress does not facilitate adaptations in social or anxiety-like behaviors

As we determined that OE of H1x in stress-naïve mice does not induce behavioral adaptations, we next tested whether the interaction between stress exposure and elevated H1x contributes to the emergence of the susceptible phenotype. Here, we applied a 10-day CSDS paradigm and SI testing to identify cohorts of susceptible and resilient mice. We then delivered HSV-H1x versus HSV-GFP to the vHipp, and completed a behavioral posttest for SI, EPM, and novelty suppressed feeding (NSF), each in distinct cohorts of mice (Fig. 4A). We again utilized the behavior pretest to behaviorally match cohorts of resilient mice receiving HSV-H1x versus HSV-GFP and behaviorally match cohorts of susceptible mice receiving HSV-H1x versus HSV-GFP, to prevent individual differences in behavior from biasing the treatment effect.

Following viral treatment in our cohorts of susceptible and resilient mice, we observed no significant difference between mice of either pretest phenotype treated with HSV-H1x versus HSV-GFP in terms of time spent interacting with the novel CD1 mouse in the SI test (Fig. 4B). We observed that viral treatment did not affect thigmotaxis behaviors, as HSV-H1x and HSV-GFP mice spent equivalent time exploring the center of an open field (Fig. 4C), nor did it affect basal locomotion as HSV-H1x and HSV-GFP mice moved a similar distance in the open field (Fig. 4D). Additionally, we saw that following CSDS, viral treatment did not alter NSF behaviors (Fig. 4F) or time spent exploring the open arm of the EPM (Fig. 4E). These results were consistent across three distinct cohorts of mice ran though this behavioral pipeline. Moreover, when these behaviors were analyzed specifically within stress-susceptible (Supp. Fig. 1) or stress-resilient (Supp. Fig. 2) mice, there remains no significant difference between HSV-H1x treated mice and HSV-GFP treated mice in any of these metrics.

We next tested whether increased H1x expression preceding the stress experience modulates responses to stress using an accelerated twice per day CSDS protocol as previously described^24,25^. Across two cohorts of mice, we did not see H1x OE induce any difference in SI or EPM behaviors compared to GFP controls following accelerated CSDS (Supp. Fig. 3).

For the time spent on the open arms of the EPM (Fig. 4E), the interaction term between test day and virus treatment met significance (p = 0.0491). This hints that the viral effect was reliant on repeated exposure to the EPM. As the pretest phenotype did not significantly contribute to variance in this test, we collapsed phenotype groups and looked at all HSV-H1x versus all HSV-GFP mice. Under these conditions, the multiple comparison between HSV-H1x versus HSV-GFP on posttest showed a trend where HSV-H1x mice explored the open arms less (p = 0.0793). This led us to question whether intra-vHipp OE of H1x impacts the capacity to recall anxiogenic spatial contexts, like the open arm of an EPM.

### Overexpression of H1x following stress does not alter fear conditioning

To test the potential impact of vHipp HSV-H1x treatment on memory and recall of anxiogenic spatial contexts, we employed the fear potentiated startle (FPS) paradigm. Here, we subjected cohorts of susceptible and resilient mice following vHipp delivery of HSV-H1x versus HSV-GFP to a fear conditioning and reversal learning paradigm according to established protocols^22^ (Fig. 5A). Following habituation to the startle chambers, we calculated baseline startle response adjusted for individual animal weight (Supp. Fig. 4). We did not find any behavior difference in startle response between HSV-H1x and HSV-GFP viral treatment, and there was no difference between susceptible and resilient mice (p > 0.05).

We subjected our mice to 10 foot shock-cue light pairings on the conditioning day and compared their proportional startle responses with and without the cue light conditioned stimulus (Fig. 5B). To confirm successful pairing of the unconditioned stimulus of the foot shock to the conditioned stimulus of the cue light, we completed a one-sample t-test comparing to a theoretical proportional startle response of 0. As both HSV-H1x and HSV-GFP mice display potentiated startle responses greater than 0, we can confirm the acquisition of a potentiated startle response (p = 0.0510, 0.0079 respectively). However, we did not observe a significant difference in potentiated startle response between groups (p > 0.05).

We also tested reversal learning following two days of extinction trials as a measure of cognitive flexibility. After the extinction trials, we again compared their proportional startle responses with and without the cue light conditioned stimulus (Fig. 5C). Using the one-sample t-test, we confirmed successful extinction of the potentiated startle response, as both groups proportional startle response returned to 0 (p > 0.05 for both HSV-H1x and HSV-GFP mice). We again observed no differences in proportional startle response between treatment groups, suggesting HSV-H1x did not alter acquisition or inhibition of conditioned fear responses in our test mice. Potentiated startle response data following conditioning and following reversal learning separated by pretest SI phenotype can be found in Supplemental Figure 5. In sum, these data and the data of Figure 4 indicate that by any of the diverse behavioral metrics utilized herein, there was no detectable behavioral difference between HSV-H1x and HSV-GFP treated mice.

## Discussion

Here we present a comprehensive behavioral analysis of the consequences of H1x OE, which is a linker histone protein that we demonstrate to be enriched within the vHipp of mice susceptible to social stress. By virally increasing H1x protein levels in the vHipp, we investigated whether it is possible to influence behavioral adaptations to stress and affect the emergence of the stress-induced susceptible phenotype. Although our viral vector did successfully increase intra-vHipp levels of H1x protein, we did not find any behavioral consequences of vHipp H1x OE, regardless of an animal’s level of stress exposure or its determined stress-related phenotype. We feel this is important to report, since there is a robust body of literature that has identified brain molecular profiles that diverge across stress-resilient and stress-susceptible mice, and have often described perturbations of these brain processes as causally controlling the emergence of stress-related behaviors^7–10,26–30^, including in the vHipp^31–37^. Despite these past insights into the brain molecular mechanisms that control stress response, it is reasonable that not all molecular profiles correlated with stress response are singularly sufficient to drive the emergence of stress-related behaviors. Here, we report an instance of correlation not leading to causation, in that elevated H1x protein levels in the vHipp were correlated with the stress-susceptible phenotype, but not causal in modulating any stress-related behaviors.

We selected HSV viral vectors, a specific neurotrophic vector^38^, for the experiments described here due to endogenous neuronal enrichment of H1x within vHipp. Single cell RNA sequencing analysis has identified that the cell types that highly express H1x are nearly exclusively neuronal, including Sox4^+^ neurons and Lhx1^+^ Cajal-Retzius neurons^39^. Both of these neuron types play active roles in normal neurogenesis and neuronal differentiation^40,41^. This is of particular interest due to multiple studies linking postnatal hippocampal neurogenesis with hippocampal performance, learning and memory tasks, and the capacity to endure stress without developing behavioral changes^42–46^. Therefore, because H1x gene expression occurs largely within vHipp neurons, we deemed targeting neurons for our H1x OE manipulation to be most appropriate for the present study.

We also explored whether the timing of vHipp H1x OE could impact behavioral responses to stress. Here, we applied CSDS before intra-vHipp viral transduction (Fig. 4). HSV expression peaks within 3-4 days and wanes by 7-10 days^38^; our post-surgery behavior tests were timed to overlap with peak viral expression. We additionally utilized an accelerated, twice per day CSDS protocol following viral transduction to test whether elevated H1x expression inherently modulates stress experience to dictate behavioral adaptations to stress (Supp. Fig. 3). As in Figure 4, we saw that HSV-H1x-treated mice did not behave differently to HSV-GFP control mice. Thus, we do not believe that H1x OE controls stress-induced behaviors regardless of whether its expression is elevated pre-versus post-stress experience.

We acknowledge the importance of considering sex as a biological variable in these types of studies. The data presented here represents experiments conducted on male mice because prior work identified H1x differential expression in the vHipp in male mice. Since we found no significant behavioral change following H1x OE in these cohorts of male mice, we elected not to expand these studies to female mice.

Previously, multiple core histone variants have been documented to play critical roles in many behavioral processes, including the fear conditioning paradigm applied in this study^28,35,37,47,48^, with particular attention being paid to histone crosstalk between different PTMs on specific histone variants influencing learning and memory^49,50^. A prominent example of this histone crosstalk involves H1x specifically altering H3 lysine methylation, in turn altering gene expression of multiple immune related genes^27,51^. Such immune genes have been well-documented to dictate many aspects of animal behavior, particularly social behaviors^52–54^. Thus, that we found the lack of behavioral consequences following H1x OE is surprising. H1 linker histones in general have previously been demonstrated to control chromatin architecture and maintain transcriptionally inactive heterochromatin states^12,13,55^, and H1x in particular induced extensive chromatin condensation in *in vitro* studies^55,56^. Indeed, other studies have discovered chemical and behavioral deviations following gene knockdown of specific histone variants^27–29,37,57^. Thus, we initially hypothesized that H1x OE in combination with stress experience would result in behavioral adaptations as a result of chromatin level and transcriptional changes within the vHipp. Understanding if other brain molecular mechanisms or organismal experience is required to enable vHipp H1x to exert casual regulation on stress-related behaviors is an important future direction of this work.

In conclusion, this study investigated whether heightened levels of vHipp H1x, a protein shown to be enriched in the vHipp of mice that develop social withdrawal behaviors following chronic stress, causally drives the development of these susceptible-like behavioral adaptations. By virally delivering H1x to the vHipp of mice, we did not identify any behavioral adaptations induced by H1x OE, implying that vHipp H1x is not singularly responsible for orchestrating stress-induced behavioral adaptations. Thus, we present a case of a biological mechanism that correlates with stress-induced behavioral deficits but is not singularly sufficient to manifest such behaviors.

## Acknowledgements

This work was supported by the NIH grants R00DA045795, P30DA033934, R01DA058958, and Blick Scholar funds to PJH, a V30 training award from the VCU MD-PhD Enrichment Fund to RKK, F31MH133309 to NLT, and F31DA057830 to JAP.

## Figures

**Supplemental Figure 1:**
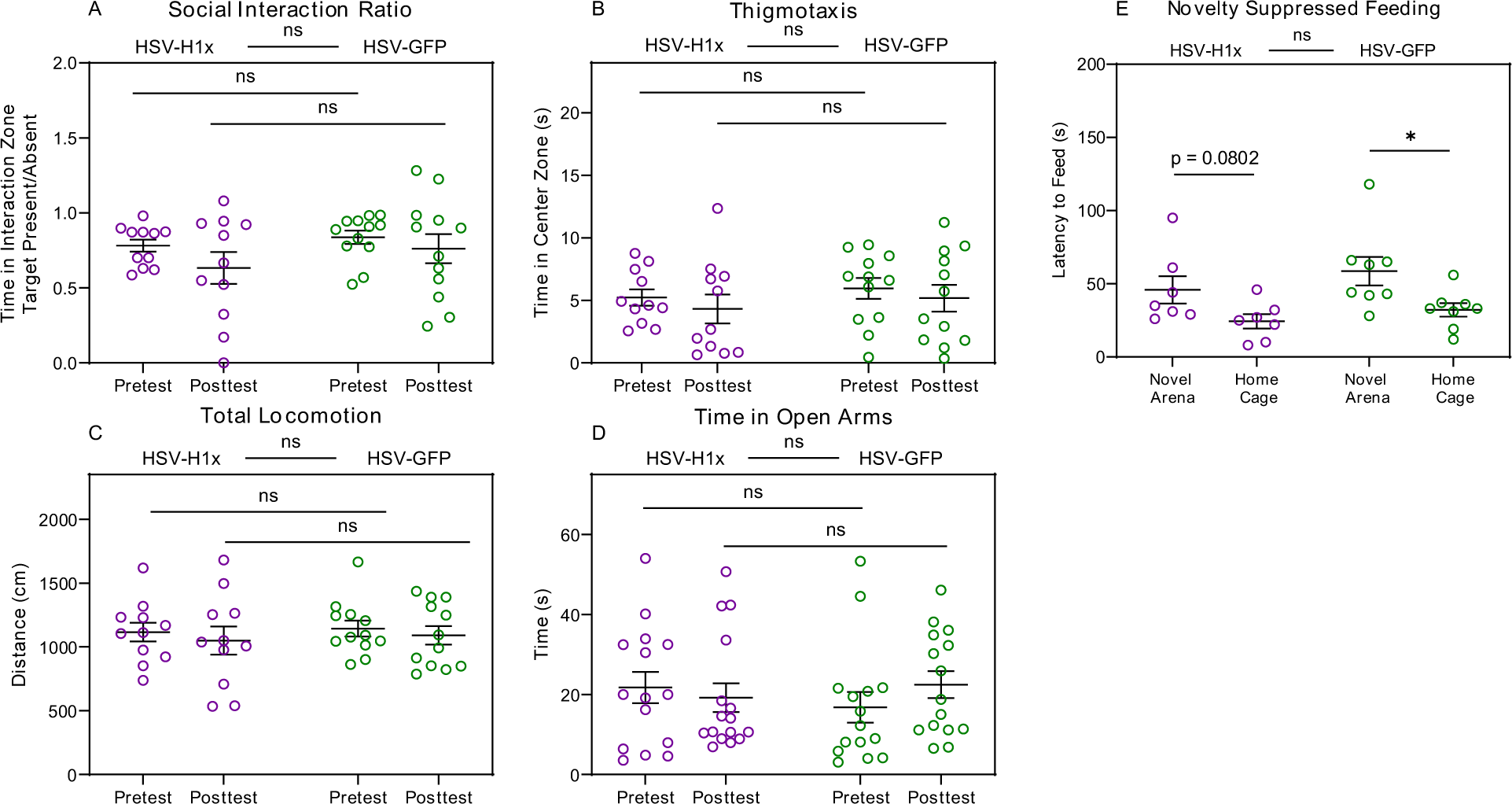
Behavioral results of intra-hippocampal overexpression of H1x in stress-susceptible mice. No change in behaviors were observed between mice treated with HSV-H1x versus HSV-GFP (Two-way ANOVA with Bonferroni’s correction, p > 0.05).

**Supplemental Figure 2:**
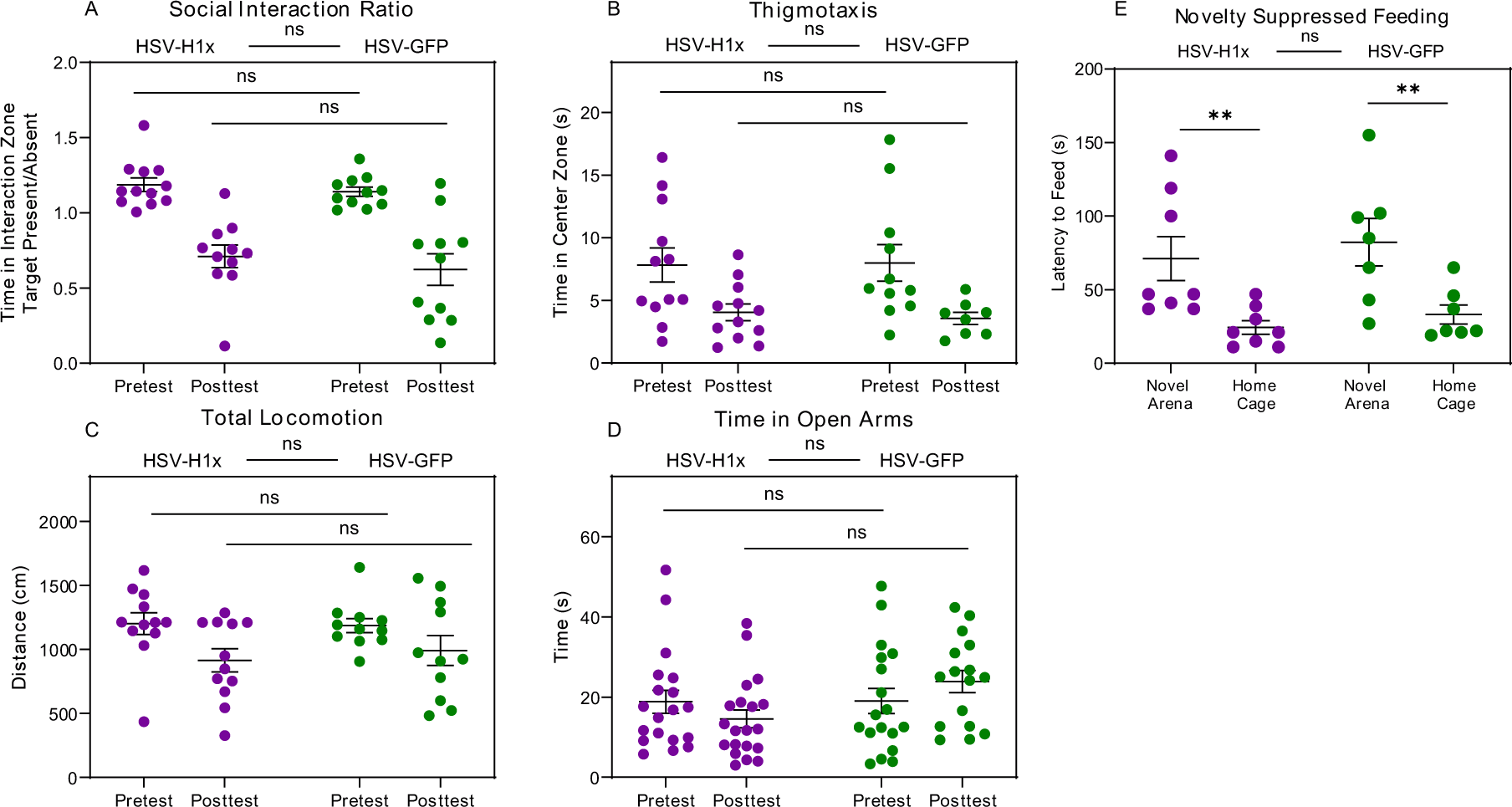
Behavioral results of intra-hippocampal overexpression of H1x in stress-resilient mice. No change in behaviors were observed between mice treated with HSV-H1x versus HSV-GFP (Two-way ANOVA with Bonferroni’s correction, p > 0.05).

**Supplemental Figure 3:**
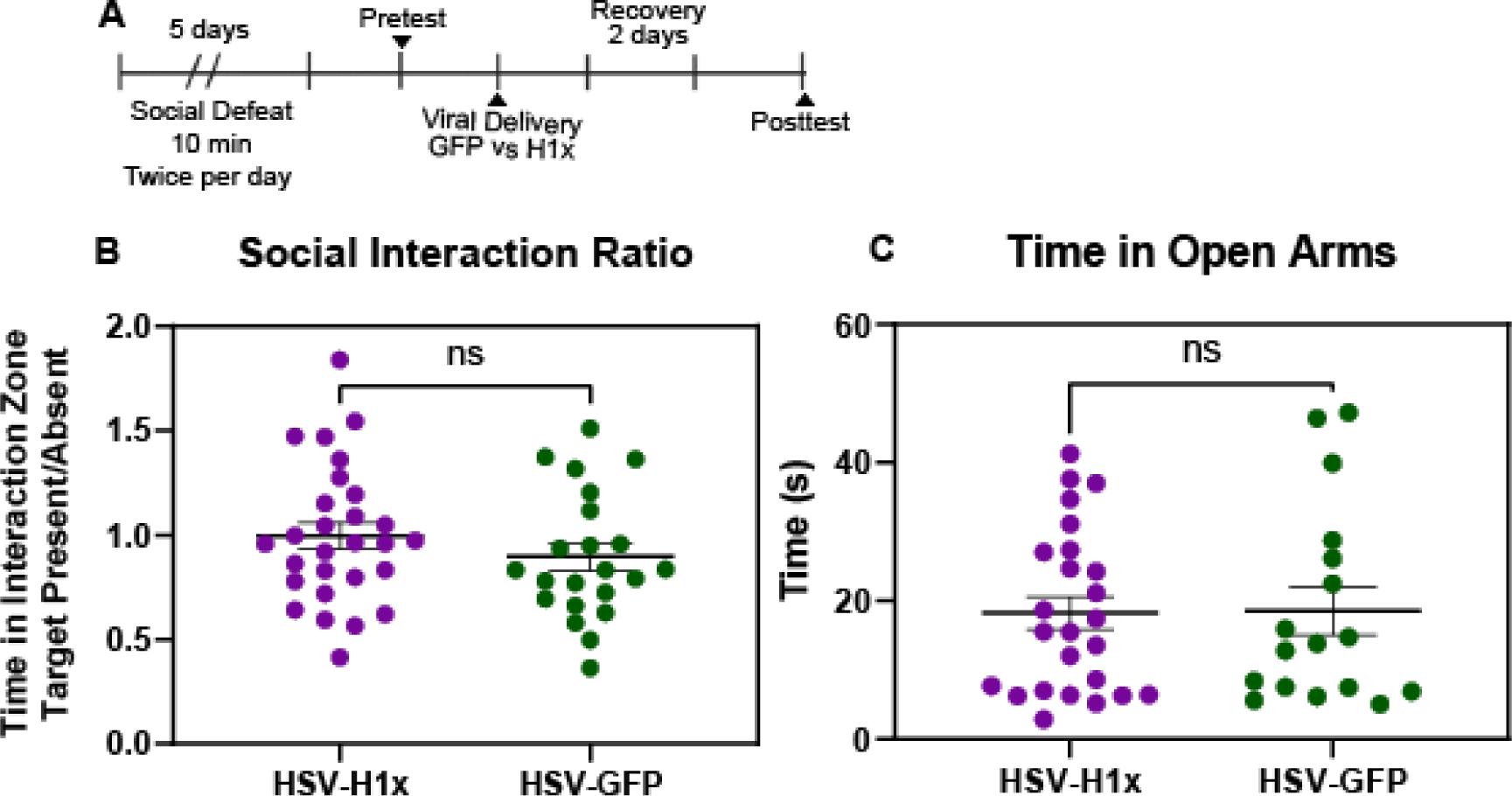
Accelerated social defeat after delivery of HSV-H1x did not alter behavioral adaptations to stress experience. (A) Experimental timeline. Accelerated twice per day social defeats were completed as previously described, with defeats spaced 6 hours apart. Behavioral testing involved social interaction and elevated plus maze behaviors. (B) No significant difference in social interaction ratios was observed between mice treated with HSV-H1x versus HSV-GFP (Two-tailed t-test, p > 0.05). Further, there was no change in the proportion of mice found to be susceptible to stress between either virus treatment group (Fisher’s exact test, p > 0.05). (C) No change in elevated plus maze exploration was observed between viral groups (Two-tailed t-test, p > 0.05).

**Supplemental Figure 4:**
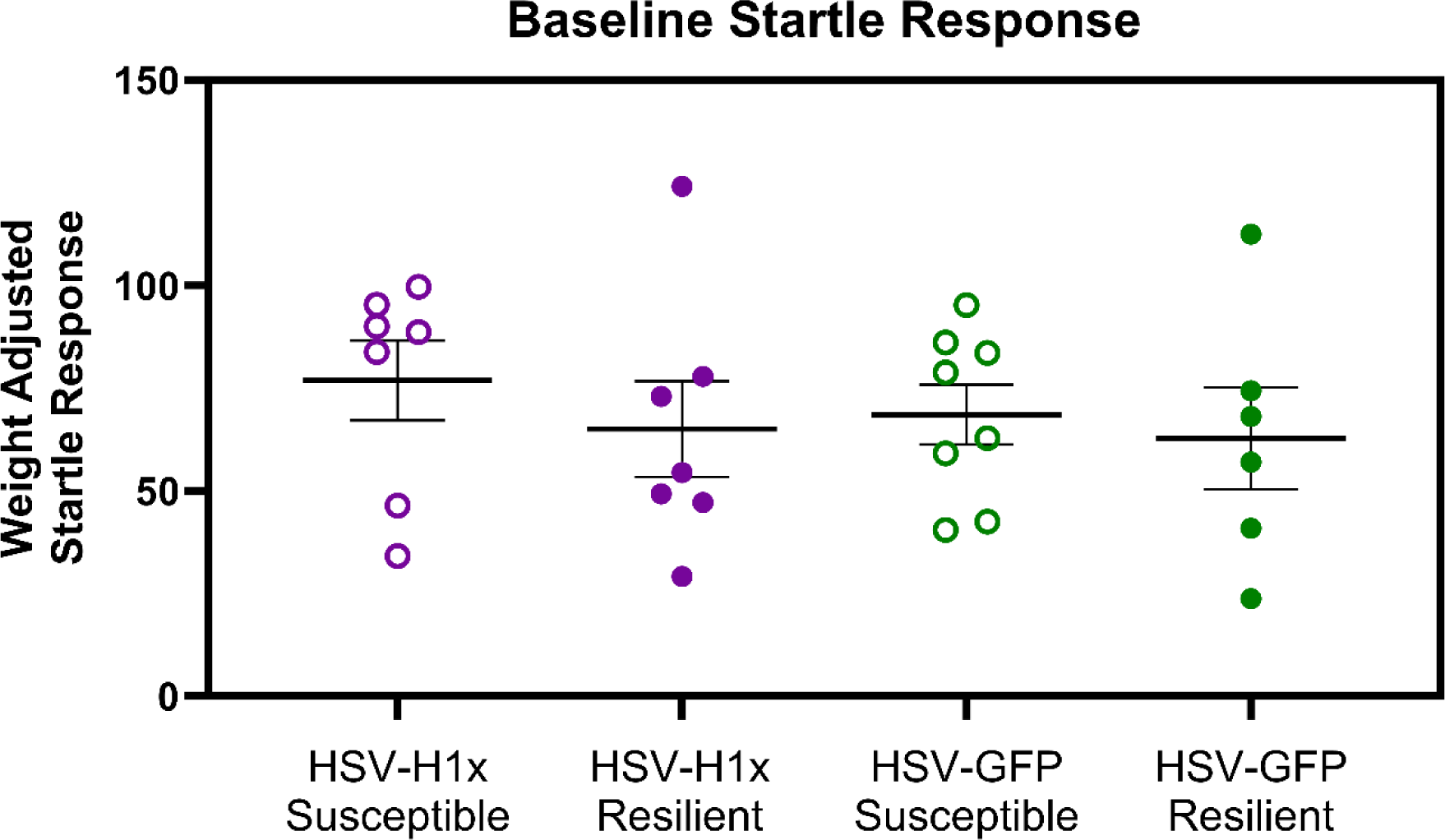
Susceptible and resilient mice treated with HSV-H1x versus HSV-GFP do not show differences in baseline acoustic startle behaviors. Following viral overexpression of H1x versus GFP, no differences in startle response to acoustic stimuli were observed between susceptible or resilient mice treated with either HSV-H1x or HSV-GFP (One-way ANOVA, p > 0.05).

**Supplemental Figure 5:**
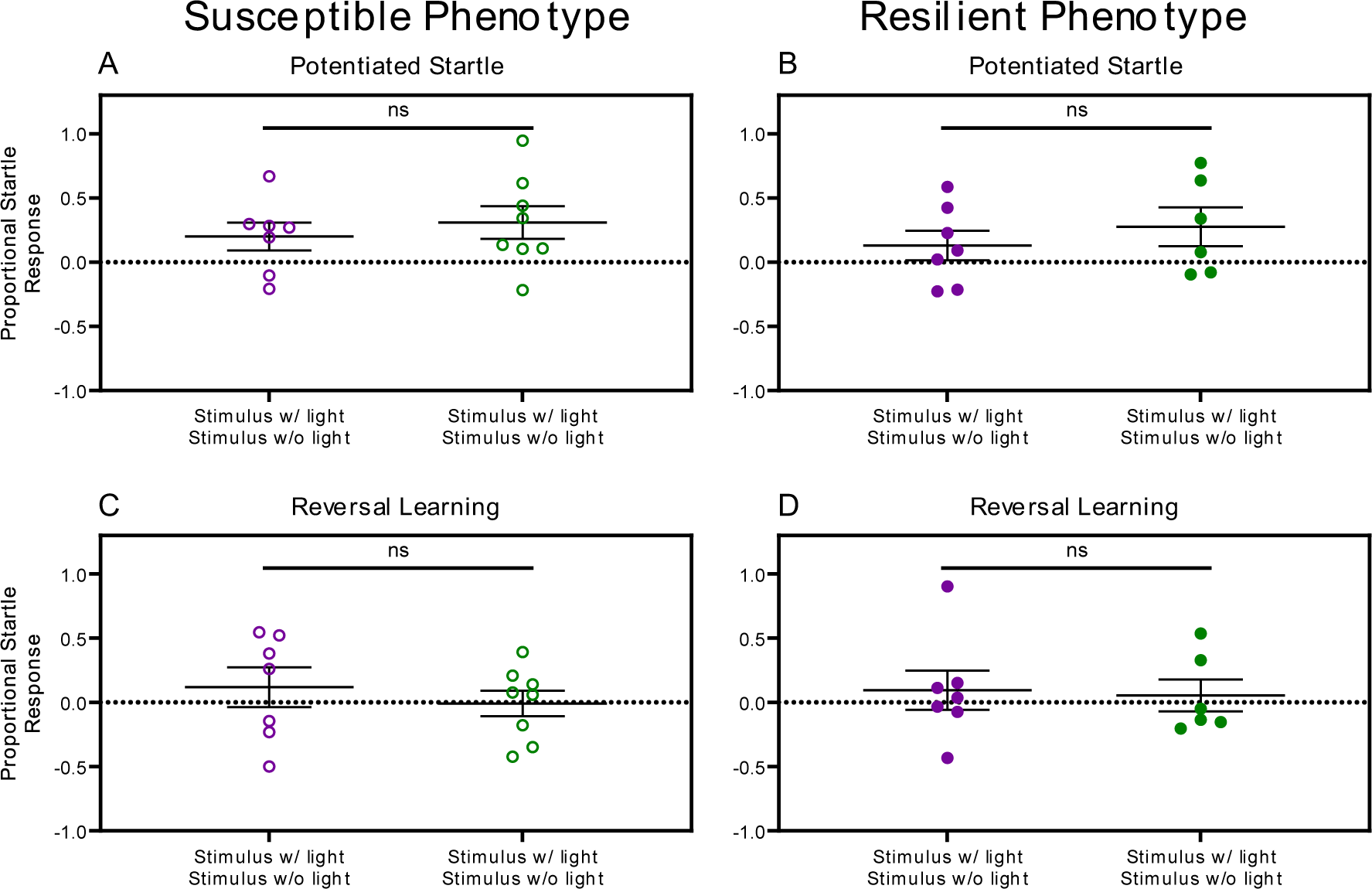
Fear-potentiated startle behaviors are not driven by a single stress-induced phenotype. Separating data by pretest phenotype does not reveal phenotype-specific deficits in either acquisition or reversal of potentiated startle (Two-way ANOVA with Bonferroni’s correction, p > 0.05).

